# Godanti Bhasma (anhydrous CaSO_4_) induces massive cytoplasmic vacuolation and cell survival response through stimulation of LC3 Associated Phagocytosis (LAP)

**DOI:** 10.1101/2020.02.01.930594

**Authors:** Subrata K. Das, Alpana Joshi, Laxmi Bisht, Neeladrisingha Das, Achariya Balkrishna, Santanu Dhara

## Abstract

Bhasmas are Ayurvedic herbo-mineral formulations that have been used since ancient times for therapeutic benefits. Godanti Bhasma (GB) is an anhydrous calcium sulfate preparation processed by heating of gypsum powder with herbal extracts. Thermo-transformation of gypsum into the anhydrous GB was confirmed by Raman and FT-IR spectroscopy. GB particle showed size range of 0.5-5 µm and neutral surface charge. Exposure to mammalian cells with GB particles showed massive vacuolation in their cytoplasm. Interestingly, no vacuolation was observed with parent gypsum particle. The result indicated that the cytoplasmic vacuolation by GB was due to its unique physicochemical property obtained during the thermo-transformation of gypsum. Using lysosomal inhibitors Bafilomycin A1 (BFA1) and Chloroquine (CQ), the process of vacuole formation was suppressed indicating GB induced vacuolation require acidic environment. The GB induced vacuolation was also found to follow dose and time dependent manner. Vacuolation often accompany with the sign of cell death whereas, in our study, massive vacuolation by GB did not induce any cell death. Moreover, GB treated cells survive with massive vacuolar process, which was reversed following post-treatment with vacuole inhibitors in GB treated cells, suggesting normal vacuolar function is essential for cell survival. Treatment of cells with GB was also found to induce translocation of LC3 protein from the nucleus to vacuolar membrane, indicating LC3 associated phagocytosis (LAP) is involved in the vacuolar process. Interestingly, the LAP function was found to be reversed in the cells treated with vacuole inhibitors. Our results provide a mechanistic correlation with GB induced vacuolation and associated LAP function, essential for cell survival.

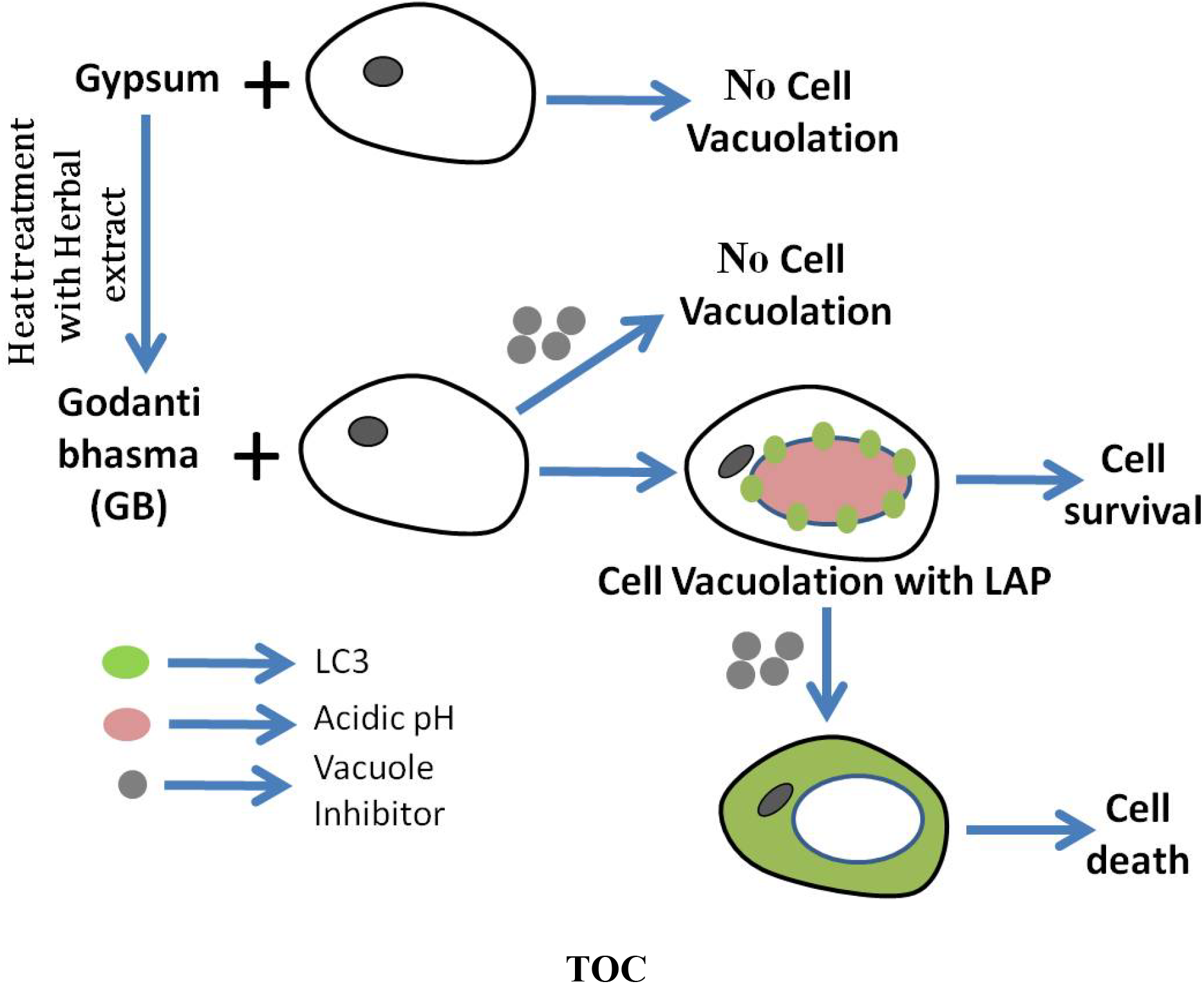

## Introduction

Eukaryotic cells develop intracellular membrane-bound organelles that compartmentalize biochemical and biophysical processes essential for cellular functions. Cytoplasmic vacuoles are one member of the organelles that serves a variety of functions including; secretory, excretory, and storage. Plant cells contain very large vacuole often occupy around 90% of total cell volume. Fungal cells also contain prominent vacuoles. In *Saccharomyces cerevisiae*, vacuole occupies approximately 25% of total cell volume. Unlike plant and fungi cells, most animal cells commonly do not contain vacuoles as regular organelles. The vacuoles in animal cells are very smaller than plant and fungi, enriched with hydrolytic enzymes called lysosome. These are about 0.2-2 μm in diameter and acidic in nature. Irreversible cytoplasmic vacuolization has been found in animal cells, when cell is infected by some viral and bacterial pathogens [1-4] as well as with the treatment of various natural and artificial low-molecular-weight compounds including medicinal drugs and industrial pollutants [5-11]. In most of the cases, irreversible cytoplasmic vacuolation is related to cell toxicity. In contrast to irreversible vacuolation, reversible vacuolation occurs as long as chemical compounds are present in medium suggesting vacuolization is a reversible process. But prolonged exposure of the chemical causes vacuole irreversible and eventually cell death [12-14].

Vacuolation is initiated by the interaction of cell membrane and particles. Researchers have tried to understand the response of the biological system during and after particle uptake in different cells and tissue model [15-17]. The mechanism of particle uptake occurs either by diffusion or endocytosis. Endocytosis is a process where extracellular or cell surface particle is internalized by the invagination as well as pinching off the plasma membrane. Usually, larger particles (>0.5 µm) internalized by phagocytosis that occurs in restricted cells (macrophage, monocyte) [18]. Phagocytosis is triggered by the interaction of the particle-bound ligand with receptors on the surface of the cell membrane; then the particle is delivered into a sealed intracellular vacuole i.e. phagosome [19-21]. V-ATPase delivered from the membranes of the endocytic pathway during phagolysosome formation is responsible for the maintenance of acidification in the vacuolar lumen [22, 23]. The maintenance of acidification in the vacuolar lumen is crucial for the hydrolytic activity of the lysosomal enzymes. Various unwanted biomolecules including proteins, nucleic acids, carbohydrates, and lipids are broken down by the hydrolytic enzymes in an acidic condition in the vacuolar lumen. Phagosomes also recruit key elements of the autophagic machinery that are responsible for conjugating LC3-family proteins to lipids at the phagosome membrane, which is called LC3 associated phagocytosis (LAP) [24]. LAP has been observed as an important mechanism to efficiently clear dying cells by professional (macrophage) and non professional phagocytes (epithelial cell and fibroblasts) [25]. The strategy of vacuole biogenesis (phagosome) in response to extracellular particle, and its further structural and functional maintenance is very important for the survival of cells. When the vacuolar function is impaired, cells become stressed and eventually cell death occurs.

Bhasmas are unique Ayurvedic medicines, prepared by incineration of metals or minerals particles processed with the herbal extract [26]. In India, therapeutic uses of these medicines are commonly evidenced since the 7^th^ century [27]. GB is a traditional ayurvedic medicine mainly used for digestive impairment [28], osteoarthritis [29] and gastric ulcer [30]. However, the mechanism of action of the drug has yet not been studied.

In the present study, we characterized the GB particle and investigated massive cytoplasmic vacuolation in murine fibroblast cell line (3T3L1) by GB particle. Here, we found that GB induced vacuolation is associated with LAP function where part of key element of autophagy machinery is involved [25]. Finally we evaluated a mechanistic correlation of vacuolar formation and associated LAP function in the survival response of cells.

## Results

### Godanti Bhasma; Anhydrous phase of gypsum

In Ayurveda, incineration procedure plays an important role in transformation of metals or minerals into therapeutic form. In the present study, GB is prepared through incineration of gypsum powder at 800°C temperature. Here we characterized the structural difference of gypsum and GB particle by using Confocal Raman in the range of 20-4000 cm^-1^, and FT-IR in the range of 600-4000 cm^-1^. In Raman spectroscopy (Fig. 1A), the presence of water in gypsum was detected by its two characteristic absorption bands at 3494 and 3406 cm^-1^ regions showing O-H stretching mode of gypsum which was disappeared in the GB (anhydrite phase). The formation of phase wise anhydrite forms of CaSO_4_-H_2_O system was confirmed by the complete disappearance of water molecules while heated at 110 to 1300°C [31]. Further, the main Raman band centered at 1008 cm^-1^ in gypsum was shifted to 1017 cm^-1^ in GB. These are υ1 symmetric stretch vibration modes of SO_4_, up-shifted following the alteration of the hydration degree. Both compounds exhibited doublet for υ2 symmetric bending of SO_4_ at 415, 494 cm^-1^ and 416, 497 cm^-1^ in gypsum and GB, respectively. The peaks at 1136 cm^-1^ in gypsum were bifurcated at 1127, 1158 cm^-1^ in GB revealed SO_4_ in υ3 asymmetric stretch vibration modes. The peaks observed at 619 and 671 cm^-1^ in gypsum and 608, 625 and 674 cm^-1^ in anhydrite gypsum are υ4 (SO_4_) asymmetric bending vibration modes. All these observations strongly indicate structural changes involving the rearrangement of sulfate ions and O-H stretching during the thermo-transition of gypsum to GB with an onset of temperature at 800 °C. The details of Raman spectra of gypsum and GB are summarized in Table S1 comparing the observed internal modes made between CaSO_4_-H_2_O system (hydrous to anhydrite phases). Some Raman spectra observed at 129, 177cm^-1^ in gypsum and 126, 168cm^-1^ in GB were attributed to Ca ion. The thermal transformation of gypsum was studied extensively by many authors [32-34].

**Fig. 1:**
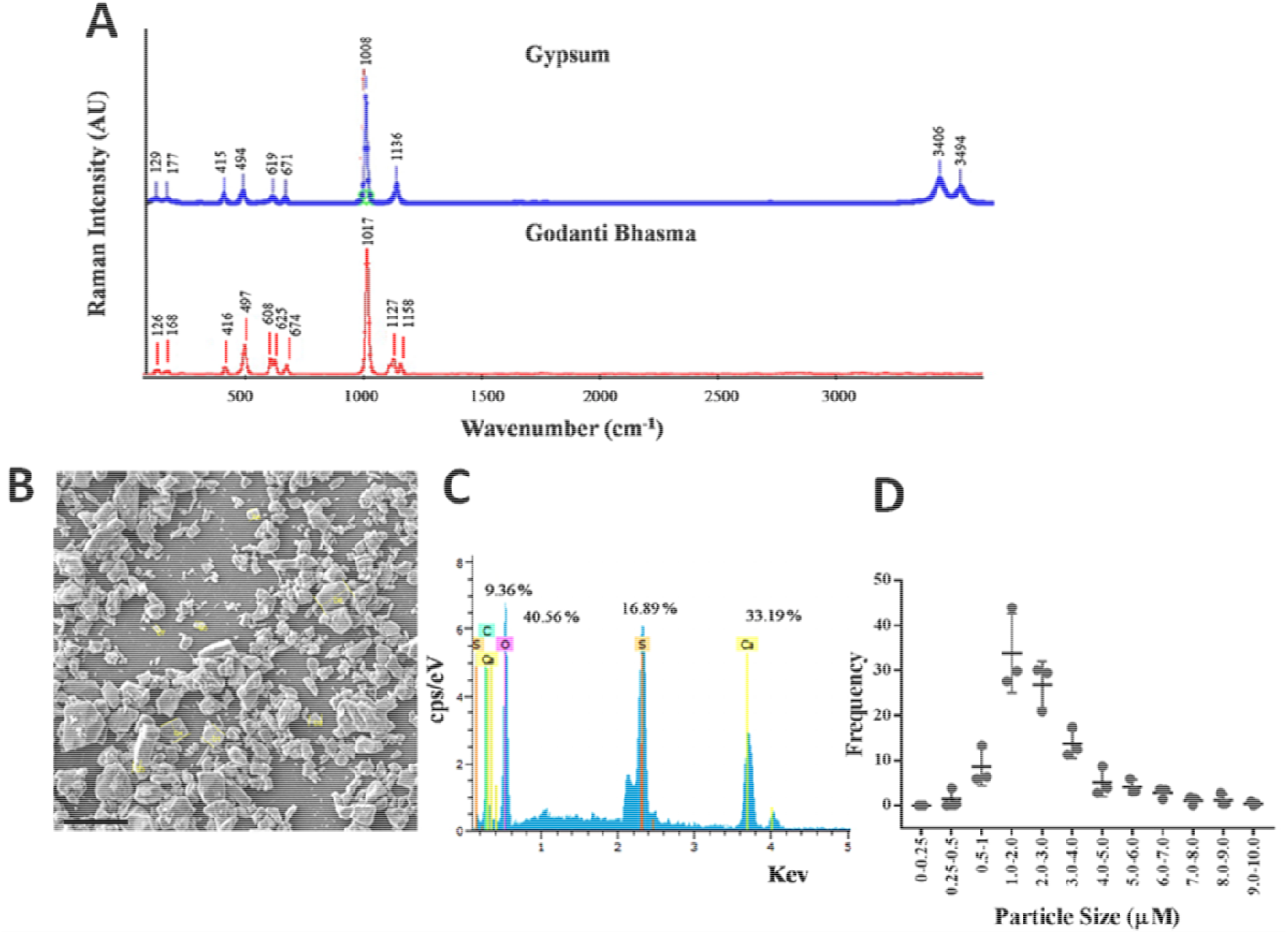
Characterization of gypsum and Godanti Bhasma. **(A)** Raman spectra of Gypsum and Godanti bhasma (GB). The Godanti bhasma is anhydrous calcium sulphate produced by thermal transformation of gypsum. Raman spectra showing two water molecules were disappeared in Godanti Bhasma. The spectral shifts were observed clearly between the two compounds. **(B)** Field Emission Scanning Electron Microscopy (FESEM) imag showing irregular particles sizes, Scale bar = 20 µm. **(C)** Compositional analysis and weight percentage (O, Ca and S) of particles calculated from Energy-dispersive X-ray spectroscopy (EDX). **(D)** Histograms showing particle size distribution based on three independent FESEM images (mean ±SEM).

The FT-IR spectrum of gypsum centered at 3527 and 3387 cm^-1^ was assigned to O-H bending vibrations, and the band at 1679 and 1617 cm^-1^ corresponds to O-H anti-symmetric stretching indicates crystalline water molecules in gypsum which were disappeared in anhydrous GB. The band at 1091 cm^-1^ in gypsum and 1097 cm^-1^ in GB associated with the stretching vibrations mode of the sulfate (SO_4_) [34] (Fig. S1).

Surface morphology and elemental composition of GB particle were analyzed by FESEM and EDX, respectively. The SEM analysis demonstrated the irregular shape with different size of GB particles ranging mean size of 0.5–5 µm (Fig. 1B and 1D). The EDX was used to detect elemental composition of Bhasma particle as calcium sulfate (Fig. 1C).

Zeta potentials were measured to evaluate electrochemical changes at the micro-particle surface due to the thermal transformation from gypsum to GB. Gypsum micro-particles showed a negative Z potential -10.42±0.98 and -11.1±0.45, whereas GB particles altered the surface charge towards neutral -1.33±2.65 and -1.75±2.41 in 10% FBS and water suspension, respectively. However, near to neutral zeta potential of particles tend to aggregate faster due to the less physical stability of the colloidal systems [35].

### Godanti bhasma induced massive vacuolation in cells

In the present study, we observed massive cytoplasmic vacuolation with various sizes of vacuole in 3T3-L1 cells exposed to the GB (Fig. 2B & S3), whereas raw gypsum powder was unable to induce any vacuolation (Fig. 2C). The vacuoles were stained with neutral red indicating their acidic nature (Fig. 2D). As pH in lysosome and vacuoles are lower than the cytoplasm, the dye penetrates inside vacuole, becomes charged and retained inside the lysosomes and vacuoles. Quantification of GB induced vacuolation was done by neutral red uptake assay in cells. A dose-dependent response was observed in 3T3-L1 cells exposed to GB with various concentrations, the vacuolation increased gradually according to the gradual increasing concentration of bhasma particles (Fig. 2G). In time course experiment, vacuolation was increased up to 24 h of bhasma treatment (Fig. 2H). It is also noted that nascent vacuoles first appeared around the perinuclear region within 2-3 hours of GB treatment, and a gradual increase in the number and size of vacuoles was observed until much of the cytoplasm occupied by a single or several vacuoles (Fig. S2). In our experiment, the bhasma particles were precipitated due to the gravitational force on the surface of the cell. However, the vacuoles were formed as long as particles are available in the culture medium. The size of vacuole approximately 1–70 μm in diameter was observed after 24 h of bhasma treatment (Fig. S3). To the best of our knowledge, extraordinarily large vacuoles were shown to induce in mammalian cells.

**Fig. 2:**
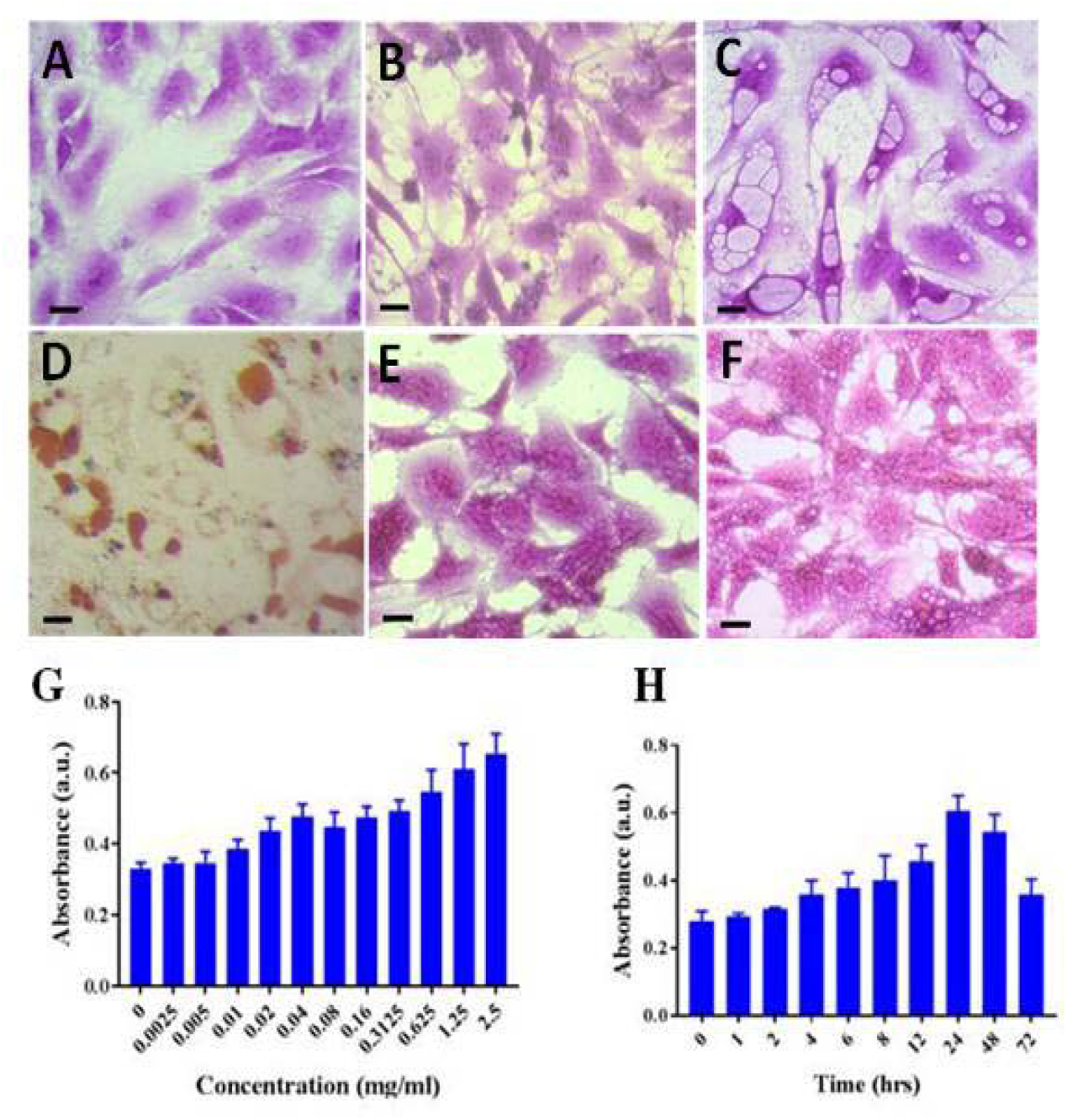
Vacuole biogenesis in 3T3-L1 cells. **(A)** Untreated cells, **(B)** Gypsum treated cells showing no vacuoles; **(C)** Godanti Bhasma treated (0.15 mg/ml) cells showing vacuolar structure in the cytoplasm of cells; **(D)** GB treated cells stained with Neutral red showing acidic pH inside the vacuole. **(E)** Co-treatment of cells with GB (0.15 mg/ml) and BFA1 (100 nM), (**F**) Co-treatment of cells with GB (0.15 mg/ml) and CQ (2 µM). Both vacuole inhibitors suppress vacuolation. Cells were stained with Crystal violet and images were captured by a bright field microscope. (Scale bar = 20 µm) **(G)** The dose response experiment showing vacuolation increased with increasing concentration of GB treatment, **(H)** GB induced vacuolation increased with increasing times upto 24 h, The results (G & H) are presented as the mean ± SD (n=6).

It was shown that GB induced vacuole formation was inhibited by Bafilomycin A1 (BFA1) indicating vacuolation require V-ATPase enzymes (Fig. 2E) that supply H^+^ to vacuole from cytoplasm. Further we found that Chloroquine (CQ) was also inhibited GB induced vacuolation (Fig. 2F). CQ is a lysosomotropic weak base; it diffuses into lysosome and vacuole where it becomes protonated and trapped. The protonated CQ then increases the vacuolar pH.

To see the specificity and degree of the vacuolation, we examined total of seven cell lines 3T3-L1, Neuro 2a, A549, MDA-MB231, MCF-7, HCT-116, and HeLa. Interestingly, all the non-phagocytic cells exhibited a similar pattern of vacuolation after 24 h of GB treatment (Fig. S 4).

### Godanti Bhasma induced vacuolation does not affect cell viability and proliferation

Initially, 3T3-L1 cells were treated with different concentrations (0-2.5 mg/ml) of GB for a period of 24 h to test its effect on cell viability. After endpoints, no significant signs of toxicity were observed in all tested concentrations of GB in 3T3-L1 cells (values over 100% should be considered as complete viability). Further, the effect was examined after the 48, 72 and 96 h of treatment on the cell viability of 3T3-L1 (Fig. 3A) and it was not found any toxicity to the cells. To explore the cytotoxic effects of GB at various concentrations (0-2.5 mg/ml) in six other cells lines (Neuro 2a, A549, MDA-MB231, MCF-7, HCT-116, and HeLa), cells were analyzed after 24 h of treatment; no toxicity was observed (Fig. 3B). Furthermore, the effects of GB on cell proliferation were investigated using *in vitro* scratch assay. The cell-free scratch area in control, as well as GB, treated 3T3-L1 cells were closed at 8 h post-scratch (Fig. 3C). This finding clearly indicated that GB does not cause any significant effect on cell viability as well as cell proliferation.

**Fig. 3:**
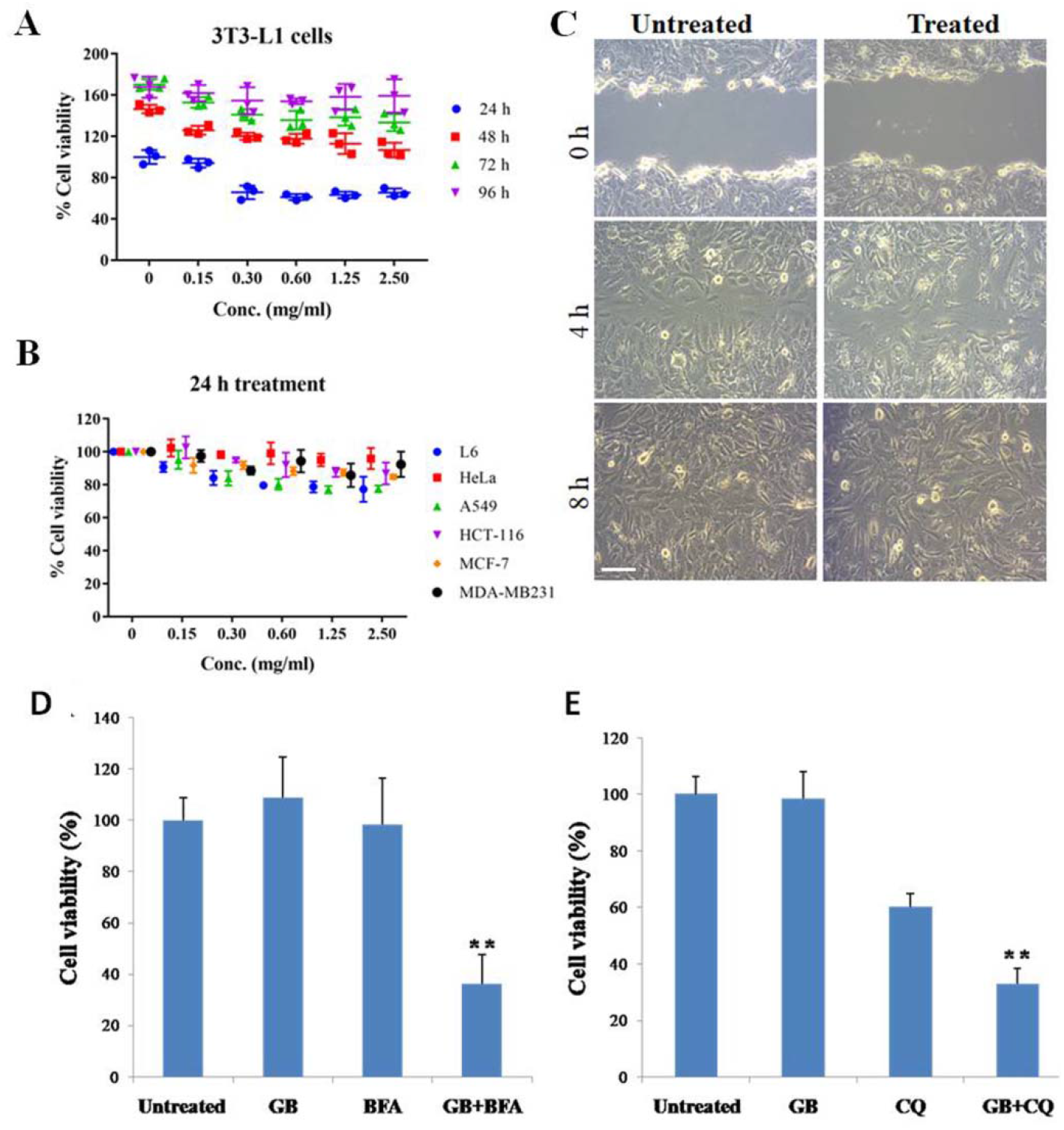
Effect of Godanti Bhasma on Cell Viability and Proliferation. **(A)** Cell viability in 3T3-L1 at different time points (24, 48, 72 and 96 h) at concentration ranges from 0-2.5 mg/ml using MTT assay **(B)** Cell viability in L6, Neuro 2a, HeLa, MDA-MB231, A549, HCT 116 and MCF-7 at concentrations vary from 0 – 2.5 mg/ml after 24 h of treatment using MTT assay, **(C)** Scratch assay in Bhasma treated 3T3L1 cell at different time points (0, 4 and 8 h). The cells were viable and proliferation was normal., Scale bar = 50 µm. **(D)** Loss of viability of GB induced vacuolated cell by post treatment of Vacuole inhibitor BFA1 with controls, and **(E)** Loss of viability of GB induced vacuolated cell by Post treatment of CQ in GB treated cells with controls. A significant toxicity of GB induced vacuolated cells was observed by the post treatment of BFA1/CQ compared to GB and BFA1/CQ controls. The concentration of GB, BFA1 and CQ were 0.3 mg/ml, 100 nM and 1 µM respectively. \*\**P*□< □ 0.01. The results are presented as the mean ± SD (n=6).

Activation of the caspase-3 pathway is a characteristic of apoptosis, and to determine the involvement of caspase-3 in GB induced vacuolation, the activation of caspase-3 by colorimetric caspase-3 assay system at different time points was examined. Exposure of 3T3-L1 cells to Bhasma particles (0.15 mg/ml) for 0, 2.5, 5, 10 and 24 h caused no increase in caspase-3 activity compared to a positive control (caspase-3). Untreated cells were used as a negative control (data not shown).

### Post treatment of vacuole inhibitors BFA1 and CQ promote cytotoxicity in GB treated cells

In our experiment, we found that GB induces massive vacuolation without any cell death. The vacuoles were reversible as vacuolar turnover after 24-72h of particle addition. To prove the vacuolar function in survival of GB induced vacuolated cells, we introduced vacuole inhibitor BFA1 (100nM) and CQ (1µM) after 24 hours of GB treatment in 3T3-L1 cells. We performed cell toxicity assay using crystal violet staining after 48 hours of the inhibitors addition, and found a significant cell death in GB+BFA1 (Fig 3D) and GB+CQ (Fig 3E) compared to respective controls. The result indicated that cell death occurs due to irreversible or defective vacuole by inhibition of vacuolar function with the post treatment of the vacuole inhibitors.

### GB induced vacuolation is associated with LAP function

LC3-associated phagocytosis (LAP) is a phenomenon distinct from autophagy, wherein LC3 translocation occurs to particle containing phagosome. To identify the LAP (LC3 lipidation), we performed immunocytochemistry with LC3A/B antibody in GB treated cells (Fig. 4A and S6). The result indicated that LC3 protein is accumulated in the nucleus of untreated cells, whereas in GB treatment it translocated to the membrane of vacuoles, suggesting activation of LAP function in GB induced vacuolated cells. We measured the fluorescence signal in whole cells and nucleus separately of GB treated and untreated using image J software (Fig. S5). The result revealed that the LC3 become down regulated in nucleus of GB treated cells compared to nucleus of untreated cells, whereas LC3 in treated cells up regulated in comparison to without treatment. Also, we estimated LC3 expression in cell lysate of 3T3L1 in different time points after GB addition compared to untreated control and CQ treatment (autophagy inhibitor) (Fig. 4B). The expression of LC3 increased (high basal level) in GB treated samples compared to untreated control where as in CQ it was highly expressed. The result indicated that the expression of LC3 is in steady state in GB treatment suggesting LAP function involved in vacuolar process. However, in CQ treatment autophagosome accumulation is enhanced due to inhibition of Autophagy flux.

**Fig. 4:**
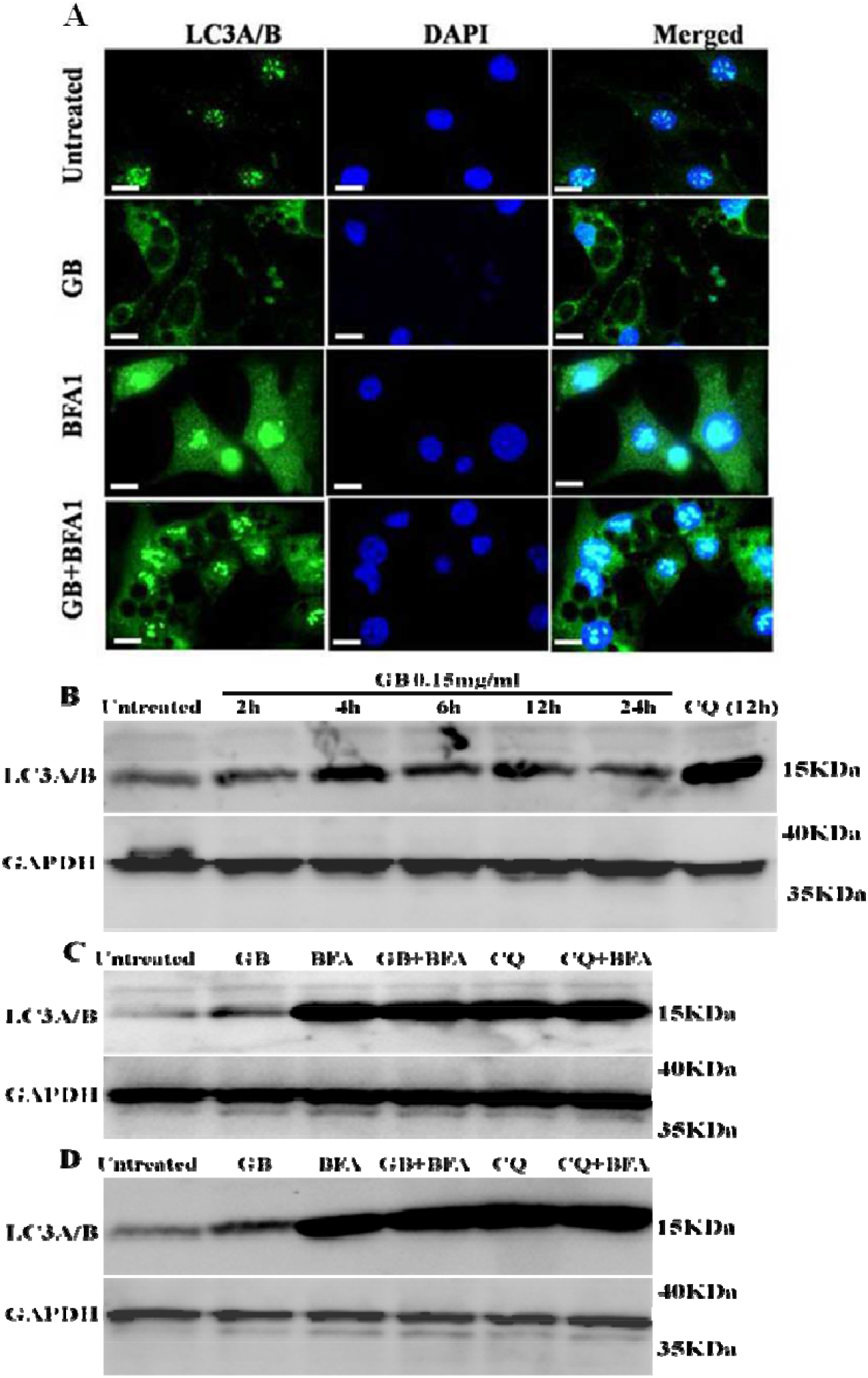
GB activated LAP like function. (A) Fluorescence image (40X) of Immuno-staining with LC3 A/B antibody showing LC3 expression in 3T3L1, LC3 expressed within nucleus in untreated cells, whereas in GB treatment, LC3 expression was found in membrane/ periphery of vacuoles indicating presence of LAP like function. LC3 expression in BFA treated cells showing LC3 expression in whole cell, and In GB+BFA, the post treatment of BFA was done in GB induced vacuolated cells, showing absence of LC3 (LC3-II) accumulation on vacuolar membrane indicating LAP like function was suppressed, whoever expression was present in whole cell. (B) The western blot showing expression of LC3 in 3T3L1 cells in different time points showing LC3 expression was in steady state compared to untreated and CQ treatment. Untreated cells was showing very less expression whereas CQ High expression, (C) Showing LC3 expression by the treatment of GB with or without autophagy inhibitors (BFA and CQ), BFA and CQ inhibited of vacuolar function as well as autophagy flux, and (D) Neuro2a experiment same as of (C). Vacuole inhibitors BFA and CQ abolished the LAP function in cells by inhibition of vacuolar function as well as autophagy flux. Scale bar = 20 µm.

To understand the relation between LAP function and vacuole inhibitor BFA1 in GB treated cells, post treatment of BFA1 (100nM) was done in GB induced vacuolated cells for 18 h. We found that the accumulation of LC3 (lipidated LC3) was suppressed on the vacuolar membrane/periphery, whereas LC3 expressed in whole cell. We examined LC3 expression with GB, BFA1 and CQ alone and GB combined with post BFA1 or CQ treatment in 3T3L1 cells by western blotting (Fig. 4C). The cell lysates were prepared after 18 h of BFA1/CQ treatment. We found that LC3 was highly expressed in presence of BFA1 and CQ with or without GB treatment. Whereas moderate LC3 expression was found in only GB treatment compared to untreated control. We also performed the same experiment in Neuro 2a cells (Fig. 4D), and found that the same expresion pattern as of 3T3L1 cells. The result indicated that LAP machanism is functional in vacuolated cells wherein it is abolished with the post treatment of vacuole inhibitors.

## Discussion

The present study sheds lights on the vacuole biogenesis induced by Godanti bhasma (GB). In this study, the most interesting phenomenon is that massive vacuolation was observed in mammalian (non-phagocytic) cells with the treatment of GB (an anhydrous CaSO_4_), whereas parent gypsum (dihydrate CaSO_4_) did not induce any vacuolation. To know this phenomenon, initially we analyzed the structural changes in gypsum and GB. The change of crystal structure on heating of gypsum was clearly evaluated by Raman and FT-IR spectroscopy. Complete removal of water molecules from dihydrate gypsum makes it more condensed crystalline form (orthorhombic anhydrite) [36] that promote their bioavailability to animal cells. According to Ayurvedic pharmaceutics, an ideal heating of inorganic minerals is essential for required physicochemical changes during Bhasma formulation [37].

Surface charge of particle is an important factor for cellular internalization. It is obvious that the interaction between positively charged particles with negatively charged cell membrane increase cellular uptake [38, 39]. It is interesting to observe from our experiments that gypsum with negative charge (z = -11) was not capable to induce vacuole formation while near neutral charged GB particle induced vacuolation effectively. Therefore, our present study reveals that charge is not an important factor for cellular internalization of GB particle.

The possible reason of such internalization of GB particles might be its structural rearrangement that facilitated to recognize cell surface receptors and, thus permeating into cells by receptor-mediated cellular uptake. Receptor-ligand interaction during phagocytosis is well studied, Fcγ *receptors* which recognize particles coated immunoglobulin *G* is the most widely studied example of phagocytosis [21]. However, few studies have been reported on the interaction of particle-bound ligand-receptors of non-phagocytic cells. The life-threatening human pathogen *Staphylococcus aureus* have an ability to internalize in a variety of non-phagocytic cells like epithelial, endothelial, fibroblast, osteoblast cells etc. The pathogen uses α5β1 integrin receptor, chaperons and the scavenger receptor CD36 to internalize into target host cells [40]. *Gratton et al* [41] revealed that HeLa cell internalized PRINT particle of 1-3 µm by several different mechanisms of endocytosis. However, the extensive study will be needed to understand the interaction between GB particle bound ligand and cell surface receptors, and downstream molecular mechanism.

After GB particle treatment, the nascent vacuoles appeared around the perinuclear region and a gradual increase in the number and size of vacuoles was observed until much of the cytoplasm occupied by a single or several vacuoles. The nascent vacuole fused with lysosome, transferring lysosomal contents to become phagolysosomes. Our experiment revealed that GB induced vacuolation required V-ATPases. V-ATPase supply high concentration of H^+^ leading higher osmotic pressure within vacuoles, and thus resulting in large vacuole due to the influx of water molecules [42-44]. V-ATPases are found on the membranes of intracellular vacuolar organelles, like endosomes, lysosomes, and phagolysosomes [45,46]. Using BFA1 as a V-ATPases blocker or using lysomotrophic agent CQ increase luminal pH that disturbs vacuolar progression in cells with GB treatment. Increasing of luminal pH of vacuole also blocks the fusion of phagosomes and late endosomes [22]. However, BFA1 and CQ induce accumulation of autophagosome in cells [47,48].

GB induced vacuolated cells was normal in proliferation which was evidenced by MTT and cell migration assay. This finding clearly indicated that GB does not cause any significant effect on cell toxicity as well as cell proliferation. Interestingly, the post treatment of BFA1 or CQ in vacuolated cells by treatment of GB showed a significant cell death compared to GB and BFA1/CQ control. In our experiment, the LAP function is activated in vacuolated cells as evidenced by accumulation of LC3-II on vacuolar membrane, whereas LC3 localized in nucleus of untreated cells. This result is strongly supported by the up regulation of LC3-II (high basal level) in GB treated cell lysate. The nuclear LC3 transported in cytoplasm as soluble LC3-I which in turn conjugated with lipid forming LC3-II and recruited to vacuolar membrane [49-51]. LAP promotes fusion of phagosome with lysosome facilitating more acidic condition in vacuolar lumen for cargo degradation [52]. The post treatment of BFA1 in vacuolated cells increase vacuolar pH and disrupts LAP function leading nonfunctional vacuolar structure. Florey et al 2015 [44] suggested that LC3 lipidation is completely blocked by BFA1 treatment. Additionally, BFA1 and CQ disrupt autophagy flux, leading accumulation of autopagosome in cells [47, 48].

The vacuolation is the sign of cell death in most of the cases by triggering apoptotic or non-apoptotic pathway [53-58]. Initially, it was believed that the accumulation of vacuoles in cell cytoplasm disrupts cellular functions and cause cell death. The mechanisms of cell death in response to vacuolation have been studied by many authors, mainly not due to vacuolation but through disruption of mitochondria, endoplasmic reticulum, Golgi apparatus, and endo-lysosomal system [8, 55, 59-61]. Moreover, the cell protects itself from a toxic substance by developing vacuole to separate from the cytoplasm. In this connection vacuolation in cells is an adaptive physiological response, presumably for damage limitation. Where damage limitation fails, cells usually die quickly. In our experiment, post treatment of BFA1 and CQ in vacuolated cells by GB showed a significant cell death compared to GB and BFA1/CQ control. Therefore, it can be concluded that the vacuole inhibitors stopped the vacuolar function by increasing pH in vacuolar lumen with non-functional LAP and thereby inhibiting further vacuolar progression. The excessive accumulation of autophagosome and defective vacuolar function is wasteful process to the cell, there by exerting cell toxicity. Whereas only GB treated cells survive with active and complete vacuolar process with a functional LAP. Therefore, our current finding indicated that LAP function is essential for survival of vacuolated cells. Moreover, cell restored its normal morphology by decreasing vacuolar volume which is associated with cellular protection from potential damaging swelling pressures.

GB is a traditional medicine used by ayurvedic practitioners in India for treating digestive impairment, acid-peptic as well as bone-related disorders. GB induced vacuole formation in mammalian cells (non-phagocytic) can be used as a powerful model to study vacuole biogenesis and further research will be needed to evaluate its pharmacological mechanism of action.

## Materials and methods

### Preparation of Godanti Bhasma

For the preparation of GB, initially, raw gypsum was coarsely powdered and washed with warm water. Then, it was suspended in sufficient quantity of lemon (*Citrus limon* L. Osbeck) juice and then subjected to moderate heat (∼80 °C) for 90 min. The obtained material was washed with warm water, dried and used for further process. In the next step, the purified powder was placed in an earthen crucible and subjected to Gaja Puta (classics nomenclature used for the quantum of heat) heating in a muffle furnace at ∼800 °C for 30 min, and then allowed for self-cooling. The obtained material was further impregnated with Aloe vera (*Aleo barbadensis* Mill.) juice and subjected to another Puta. Finally, white colored Godanti Bhasma is obtained [31].

### Characterization of raw gypsum and Godanti Bhasma (GB)

In this study, initially Confocal Raman (WITec Confocal Raman system; model: Alpha300 series), and Fourier-transform infrared (FT-IR) spectroscopic techniques were used to characterize the changes in raw gypsum and GB. Raman images and spectra were recorded using ultra-high-throughput *spectrometer* (UHTS) equipped with a charge-coupled device (CCD) camera, diode laser used for 532 nm excitation, and microscope (Zeiss 100x air objective with numerical aperture 0.9). The FT-IR measurements were obtained at room temperature using AGILENT spectrometer (model: CARY630) equipped with ATR cell attached with Micro Lab PC software. The data was recorded three times from three different set of samples for both experiments.

Further, the surface morphology and particle size distribution of GB sample were observed by Field emission scanning electron microscopy (FESEM) (TESCAN; model: MIRA3) technique. For this sample was anchored on the sample holder, and morphology was probed on selected points to determine elements with the help of detector inbuilt with energy dispersive X-ray analyzer (EDX) (Rigaku; model: XFlash 6I10) at 0-10 keV. Total three different set of particles were examined during scanning. Histograms of particle size distribution were made on three independent FESEM images.

Zeta potentials were measured to evaluate electrochemical changes at the microparticle surface due to the thermal transformation from gypsum to GB. For this samples were suspended in distilled water as well as 10% FBS and surface charged were assessed by zeta sizer nano (Malvern Panalytical, UK, ZS90). Five samples of each set were recorded.

### Cell Culture

Dulbecco’s modified Eagle medium (DMEM), fetal bovine serum (FBS), Antibiotics, Trypsin-EDTA solution were obtained from Thermo Fisher, MA USA. Cell lines (3T3-L1, L6, Neuro 2a, HeLa, HCT-116, A549, MDA-MB231, and MCF7) were procured from National Centre for Cell Sciences (NCCS), Pune, India. The cell was cultured in DMEM (GIBCO BRL, Grand Island, NY, USA) supplemented with 10% inactivated Fetal Bovine Serum (FBS), penicillin (100 IU/ml), streptomycin (100 µg/ml) and amphotericin B (5 µg/ml) in a humidified atmosphere of 5% CO_2_ at 37 °C. The cells were dissociated with Trypsin-EDTA solution. The stock cultures were grown in 25 cm^2^ culture flasks, and experiments were carried out in 96- and 6-well plates (Tarsons India Pvt. Ltd., Kolkata, India).

For cell culture experiment, GB powder (100 mg) was suspended with complete DMEM media (1 ml supplemented with 10% FBS) by vortexing, and leave the tube on a stand for 1 min to settle down larger particle, serial dilution was done using 500 µl of GB suspension. All cell culture experiment (except dose-response experiment) was conducted using 5^th^ dilution of GB suspension.

### MTT assay

Cell viability was determined using MTT assay in 3T3L1. Cells were seeded (7500 cells/well) in 96-well plate and incubated for 24 h. After 24, 48 and 72 h of incubation in presence of GB with serial dilutions (0 – 2.5 mg/ml), the culture medium of each well with or without extract was removed completely from the assay plates and replaced by fresh culture medium (100 µL). MTT (Thiazolyl Blue Tetrazolium Bromide) solution (10 µL of 5 mg/mL), (Thermo Fisher, MA USA) was added into each well to achieve a final concentration of 0.45 mg/mL before incubated for 3 h at 37 °C. After 3 h, the culture medium with MTT was carefully removed followed by addition DMSO (100 µL) (Himedia, India) to dissolve formazan crystals, and then incubated for 1 h before recording the optical density (Envision plate reader, California, USA) at 540 nm. Cell viability test was also performed in six different cell lines (Neuro 2a, A549, MDA-MB231, MCF-7, HCT-116, and HeLa) at 24 h of culture. The results are presented as the mean ± SD (n=6).

### Cell migration assay

Cells were seeded in a 6-well plate (0.5×10^6^ cells/well), incubated up to 100% confluent. Cells were treated with and without GB. The monolayer of cells was scratched (3 scratches) with a pipette tip after 12 h of treatment, and cells were imaged at 0, 4, 8h post-scratch. The cell-free areas per treatment group were used for analysis.

### Caspase assay

The Caspase-3 colorimetric assay was also conducted according to manufacturer instructions (Sigma-Aldrich, MO, USA). 3T3-L1 cells were treated with Bhasma for 0, 2.5, 5, 10, and 24 h. The concentration of the p-nitroanilide (pNA) released from the substrate was calculated from the absorbance values at 405 nm. Three samples were done for each time points.

### Crystal Violet staining

Crystal Violet (High Media, India) staining in cells was done for microscopic imaging. Treated cells were washed with 1X PBS, fixed with formaldehyde (10%) for 15 min. After fixation, cells were washed with water and stained with 0.5% (w/v) crystal violet (25% (v/v) methanol) for 25 min, subsequent washing the cells with water until no color was eluted, and images were taken by Bright Field microscope (PrimoVert. Zeiss, Germany).

### Neutral Red staining

Relative vacuolation was quantified based on the uptake of the Neutral Red dye (High Media, India) in mammalian cells (Kannan et al. 2014). The experiments were performed in 96-well plates, neutral red (100 µl of 0.5 mg/ml) was added in each well, incubated for 4 h in CO_2_ incubator, washed 3 times with PBS, eluted the neutral red by adding destining solution (100 µl containing 50% dehydrated ethanol, 49% deionized water and 1% glacial acetic acid). Neutral Red uptake was determined using a microtiter plate reader (EnVision Multimode plate reader, Perkin Elmer, USA) to measure the absorbance at 540 nm.

### Immunocytochemistry

After treatment, cells were washed with PBS (2 times) and fixed with 4% paraformaldehyde for 15 min. Following three further PBS washes, cells were permeabilized with 0.2% Triton in PBS for 10 min. Cells were blocked in blocking buffer (1% BSA, PBST) for 1h at room temperature and incubated with primary antibody LC3A/B (Thermo Scientific, USA) overnight at 4 °C. Cells were again washed with PBS (3 times, 10 min each), incubated with secondary antibody Goat Anti Rabbit IgG H&L dylight 488 (Thermo Scientific, USA) for 1 h, and then followed by PBS wash (3 times, 10 min each). Only secondary antibody (no primary ab) in treated cells was used as a control. Cells were treated with DAPI (3 g/ml) in PBS for 5 min, wash again with PBS (3 times, 10 min each), Slides were mounted with 50% glycerol. Microscopy was done with fluorescence microscope (Mantra, Perkin Elmer, USA). The immunocytochemistry was done three times.

### Western blotting

Cells were washed with PBS, scraped into ice-cold RIPA (150 mM NaCl, 50 mM Tris-HCl, 1% Triton-X-100, 0.1% SDS, 0.1% sodium deoxycholate) buffer and lysed for 10 min on ice. Lysates were centrifuged for 12 min at 4°C. Supernatants were then separated on 15% polyacrylamide SDS-PAGE gels and transferred to a PVDF membrane. The membrane was blocked in TBST (50 mM Tris-Cl, pH 7.6, 150 mM NaCl, 1% Tween 20) + 3% BSA and incubated overnight at 4°C with primary antibody LC3A/B diluted in blocking buffer. Blots were incubated with HRP conjugated to secondary antibody (Thermo Scientific, USA) and protein detected using enhanced chemiluminescence (Thermo Scientific, USA). The blot was striped with striping buffer (0.1M glycine 0.1% SDS 1% Tween20, pH to 2.2) and reproved with GAPDH antibody.

## Acknowledgment

We are thankful to Patanjali Research Foundation Trust, Haridwar, India for financial support. The authors thankful to Dr. Kiran Ambatipudi and Prof. P. Roy, Dept. of Biotechnology, IIT Roorkee, for providing lab facility for conducting experiments. We are also thankful to the Central Building Research Institute (CBRI) under the Council of Scientific and Industrial Research, India for FESEM-EDX analysis. We thank to Dr. Niti Sharma for helping FTIR of particle, Deepika Mehra for helping in immunocytochemistry cells and Vaidya Vanamra Sharma for writing method of GB formulation. We would like thank to Toshniwal Brothers (SR) Pvt. Ltd., India for providing the facility of Raman spectroscopy. We thank Mr. Nantu Dogra, SMST, IIT-Kharagpur for repeating FTIR and few cell culture experiment to check reproducibility.

## Author contribution

S.K.D. conceptualization, conducted Particle characterization, cell culture experiments, immunostaining, western blot, analyzed all data, manuscript writing, reviewing and supervised overall studies; A.J. conducted the immunocytochemistry, analyzed the data and manuscript writing; L.B. assisted cell experiment, cell staining procedure and western blot; N.D. performed particle charge; A.B. helped for resource; S.D. checked all data, reproducibility and review manuscript.

## Supporting Information

Supplemental Figure 1: Comparison of FT-IR spectrum of Gypsum and Godanti Bhasma. Figure 2: Cell images showing changes of vacuole morphology during vacuole biogenesis at different time points. Figure 3: Cells showing different sizes of vacuoles in GB treatment. Figure 4: GB induced Vacuolation in different cell lines. Figure 5: Comparison of intracellular fluorescent signals quantification whole cell vs. nucleus. Figure 6: LC3 expression in L6 cells. Supplemental Table 1: Comparison of Raman Spectra of Gypsum and Godanti Bhasma.

