## Supplementary Figure and legends for "Godanti Bhasma (anhydrous CaSO_4_) induces massive cytoplasmic vacuolation and cell survival response through stimulation of LC3 Associated Phagocytosis (LAP)"

Author’s Affiliation:

1- Drug Discovery and Development Division, Patanjali Research Institute, Patanjali Research Foundation Trust, NH-58, Haridwar-249405, Uttarakhand, India.

2- School of Medical Science and technology, IIT Kharagpur-721302, India.

3- Shobhit Institute of Engineering & Technology (Deemed-to-be-University), Meerut-250110, India.

4- Department of Biotechnology, IIT-Roorkee, Roorkee-247667, India.

\# Both authors made equal contributions

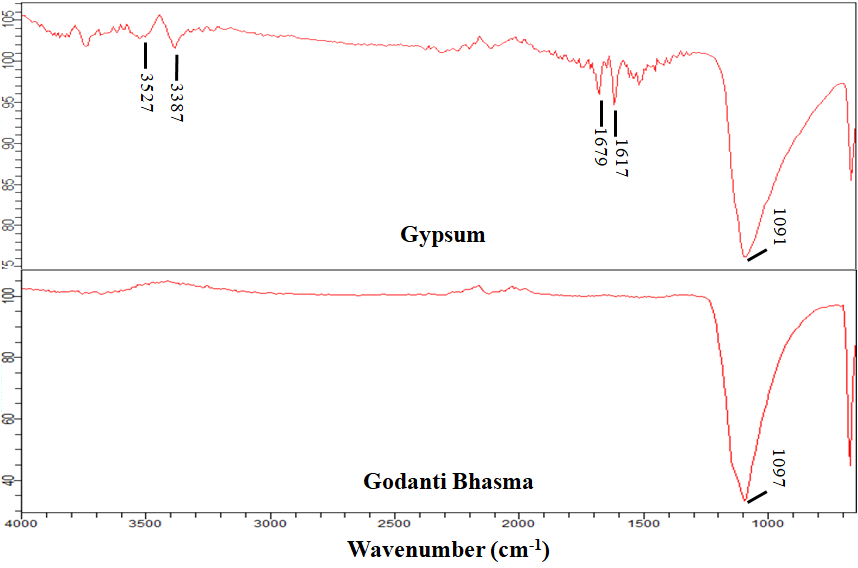

### Fig. S1: FT-IR spectrum of Gypsum and Godanti Bhasma. The spectra show water molecules disappeared in GB due to the thermal transformation of gypsum.

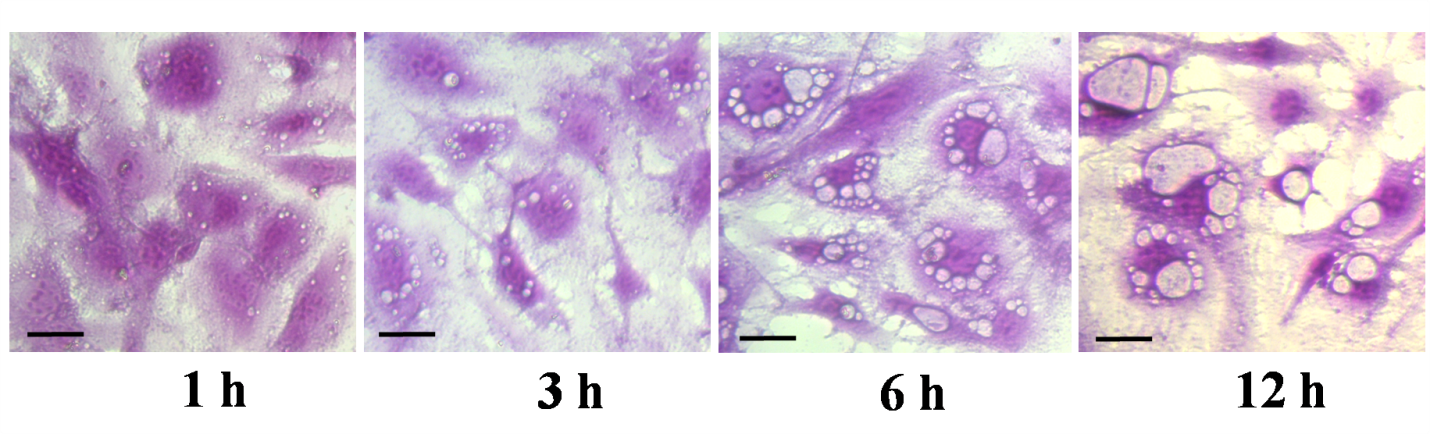

**Fig. S2:** Crystal violet stained cell images showing changes of vacuole morphology during vacuole biogenesis at different time points. Scale bar = 20 µm.

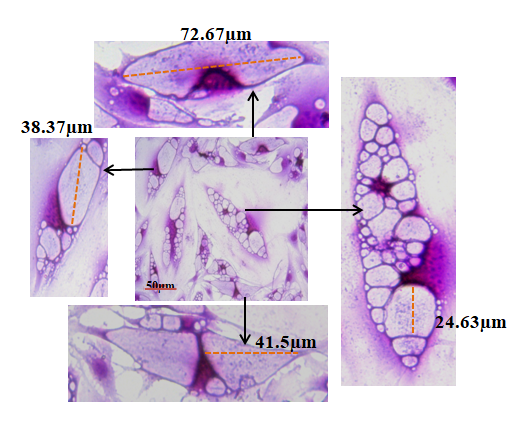

**Fig. S3:** Crystal violet stained cells showing different sizes (1-70 µm) of vacuoles in GB treated cells.

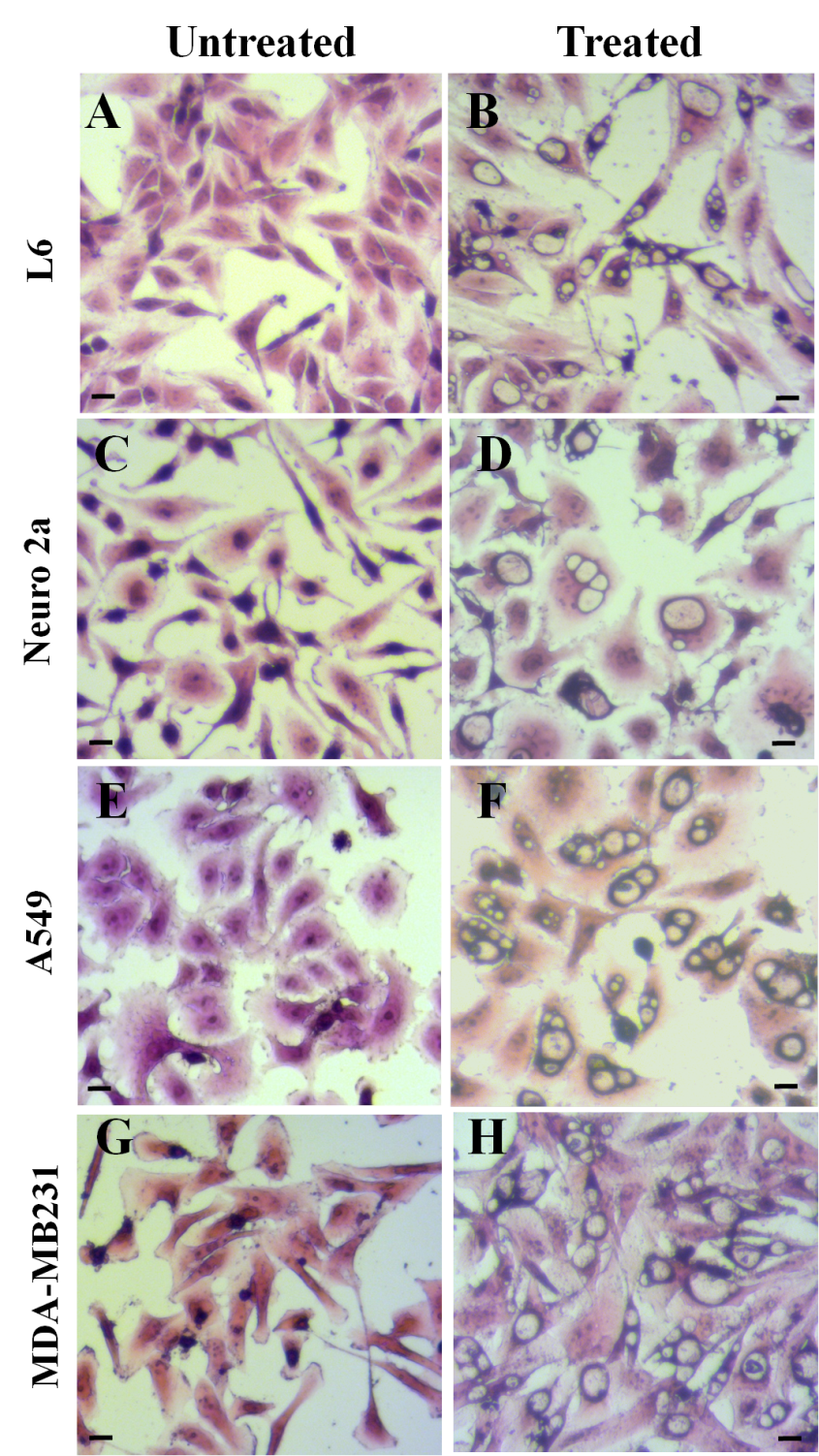

**Fig S4:** GB induced Vacuolation in different cell lines (L6, A549, Neuro 2a and MDA-MB231). Untreated (A, C, E, G) and GB treated (B, D, F, H). Vacuoles are marked with arrow (yellow colored). Scale bar = 20 µm.

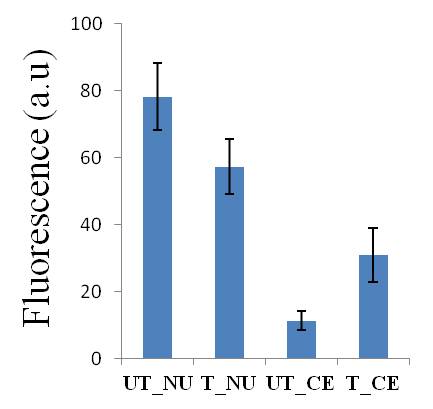

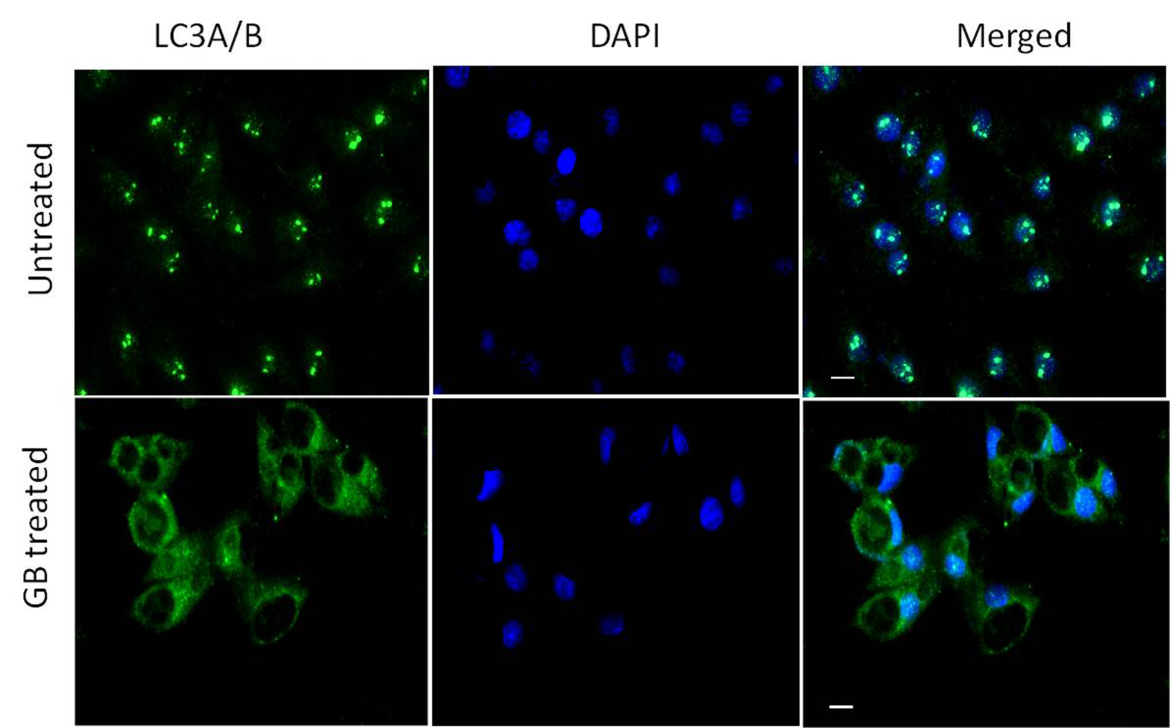
**Fig-S5** Intracellular fluorescent signals of 3T3L1 cells (untreated and GB treated) were quantified using the ImageJ software (20 cells analyzed). Quantification was done total cell vs. Nucleus and comparison was plotted.

**Fig. S6: LC3 expression in L6 cells.** (A) Fluorescence image (40X) of Immuno-staining with LC3 A/B antibody showing LC3 expression in L6 cells, LC3 expressed within nucleus in untreated cells, whereas in GB treatment, LC3 expression was found in membrane/ periphery of vacuoles indicating presence of LAP like function. Scale bar = 20 µm.

**Table S1 Raman Spectra of Gypsum and Godanti Bhasma (GB) in the CaSO_4-_H_2_O System**

| Material | ν_2_ (SO_4_) | | ν_4_ (SO_4_) | | | ν_1_ (SO_4_) | ν_3_ (SO_4_) | | (H_2_O) | |
| --- | --- | --- | --- | --- | --- | --- | --- | --- | --- | --- |
| Gypsum | 415 | 494 | 619 | 671 |  | 1008 | 1136 |  | 3406 | 3494 |
| GB | 416 | 497 | 608 | 625 | 674 | 1016 | 1127 | 1158 |  |  |
